# Structural and functional characterisation of a reconstructed ancestral strigolactone receptor

**DOI:** 10.1101/2025.08.03.668049

**Authors:** Andrew J. Tuckey, Andrew C. Marshall, Emel Rothzerg, Tom Bennett, Charles S. Bond, Mark T. Waters

**Author notes:** The authors declare no conflicts of interest. Raw data and biological material will be made available upon reasonable request to the corresponding author (MTW).

## Abstract

Strigolactones are phytohormones that regulate shoot branching and promote root interactions with arbuscular mycorrhizal fungi. In seed plants, strigolactone perception begins with the enzyme-receptor DWARF14 (D14), an α/β-hydrolase that is believed to have evolved via gene duplication from the karrikin receptor, KARRIKIN INSENSITIVE2 (KAI2). D14 and KAI2 are highly homologous at the sequence and structural levels, and both proteins bind and cleave similar butenolide compounds. Nevertheless, the two signalling pathways are distinct, as the activity of one receptor cannot compensate for the loss of the other. Here, we apply ancestral sequence reconstruction to generate a D14 protein representative of seed plants to study the evolution of substrate specificity, and to explore desirable traits for protein engineering. We describe the structure, as well as the *in vitro* and *in planta* activity of ancestral D14, showing that substrate specificity does not meaningfully differ from that of *Arabidopsis thaliana* D14. We also demonstrate that ancestral D14 shows higher recombinant yields, greatly increased thermostability, and enhanced catalytic activity relative to D14 from *A. thaliana*. This work provides insight into the evolution of phytohormone signalling and presents a robust scaffold for the application of D14-type proteins in synthetic biology.

**SIGNIFICANCE STATEMENT:** Ancestral sequence reconstruction of the strigolactone receptor DWARF14 (D14) reveals that its substrate specificity has remained largely conserved in seed plants. Additionally, the ancestral reconstruction exhibits superior recombinant yields, thermostability, and catalytic activity relative to *Arabidopsis thaliana* D14, providing both evolutionary insights and promising utility for synthetic biology applications.

## INTRODUCTION

Strigolactones (SLs) are a class of butenolide phytohormones that regulate numerous processes in land plants. SLs were initially identified as molecules exuded into the soil from plant roots that stimulate the germination of parasitic weeds in the genus *Striga* (Cook et al., 1966). The benefit of SLs for the host plant remained enigmatic until it was observed that the phytohormones also promote symbiosis with arbuscular mycorrhizal (AM) fungi (Akiyama et al., 2005). SLs additionally control shoot architecture via the inhibition of shoot branching (Gomez-Roldan et al., 2008; Umehara et al., 2008), offering a promising avenue for enhancing yields in crop plants (Kelly et al., 2023).

DWARF14 (D14), an α/β-hydrolase, assumes a dual role as both an enzyme and receptor for SLs (Hamiaux et al., 2012; Waters et al., 2012; de Saint Germain et al., 2016; Yao et al., 2016). The exact mode of action for SL signalling through D14 is incompletely resolved. The SL undergoes nucleophilic attack upon the butenolide D-ring by the catalytic serine residue, is cleaved, and is subsequently released from the protein (Hamiaux et al., 2012), but it is unclear which specific steps are necessary for signal transduction. One model posits that the hydrolysis of the D-ring bridges the catalytic serine and histidine to form a covalently linked intermediate molecule, which is a required conformation for signal transduction (de Saint Germain et al., 2016; Yao et al., 2016). This model has been disputed due to inconsistency between the time taken to hydrolyse SLs being much longer than the downstream effects of D14 signalling (Zhao et al., 2013; Seto et al., 2019). Additionally, a mutant protein lacking a catalytic aspartic acid residue can facilitate signalling, despite being unable to hydrolyse SLs (Seto et al., 2019). This result implies that binding alone is sufficient for signal transduction, with hydrolysis occurring afterwards. Nevertheless, this binding-sufficient model has itself been challenged, as it explains neither the invariant conservation of the aspartic acid residue in nature, nor the observed hydrolysis of non-bioactive butenolides (Yao and Waters, 2020).

Regardless of the precise mode of action, upon interaction with SLs, D14 undergoes a conformational shift that facilitates a protein-protein interaction with MORE AXILLARY GROWTH 2 (MAX2), an F-box protein that forms part of a SCF (SKP1–CULLIN–F-BOX) E3 ubiquitin ligase complex (Hamiaux et al., 2012; Yao et al., 2016; Shabek et al., 2018). In such complexes, the F-box protein specifies protein targets for polyubiquitination, culminating in their subsequent degradation via the 26S proteasome (Woo et al., 2001). The specific targets for D14-mediated signalling are SUPPRESSOR OF MAX2-LIKE 6/7/8 (SMXL6/7/8) in *Arabidopsis thaliana* (Soundappan et al., 2015), orthologous to DWARF53 in *Oryza sativa* (Jiang et al., 2013; Zhou et al., 2013). Notably, the D14-MAX2 complex has also been shown to interact with and mediate the proteolytic degradation of paralogous proteins SMAX1 and SMXL2 to facilitate the regulation of hypocotyl elongation (Wang et al., 2020) and osmotic stress tolerance (Li et al., 2022).

SMAX1 and SMXL2 are the downstream targets of KARRIKIN INSENSITIVE2 (KAI2) signalling (Stanga et al., 2013; Soundappan et al., 2015; Stanga et al., 2016; Khosla et al., 2020). KAI2 defines a parallel pathway involved in the perception of abiotic butenolides, known as karrikins, that are derived from smoke (Waters et al., 2012). In part because karrikins are far from ubiquitous in the environment, KAI2 signalling is thought to mediate perception and response to a butenolide phytohormone (“KAI2 ligand”, or KL) that is distinct from SLs (Conn and Nelson, 2016; Yao et al., 2021; Waters and Nelson, 2023). As *KAI2* homologues are found in all land plants and charophyte algae but *D14* homologues are limited to seed plants, it is thought that D14-mediated signalling evolved from duplication and neofunctionalisation of KAI2, concomitant with diversification of the SMXL family (Bythell-Douglas et al., 2017; Walker et al., 2019).

As homologous α/β-hydrolases involved in the perception of butenolide phytohormones, KAI2 and D14 show overlapping substrate preferences. Both respond to the synthetic SL *rac*-GR24, but they have opposing stereoselectivity. Over a range of species, KAI2 prefers the GR24 enantiomer with a 2ʹ*S* D-ring (GR24*^ent^*^−5DS^) over a 2ʹ*R* D-ring (GR24^5DS^) (Scaffidi et al., 2014; Carbonnel et al., 2020; Sun et al., 2020; Meng et al., 2021; Mizuno et al., 2021; Guercio et al., 2022; Kodama et al., 2022). In contrast, all known natural SLs have stereochemistry comparable to the latter, and these enantiomers are most bioactive through D14 (Umehara et al., 2015; Flematti et al., 2016). Another differing facet of ligand specificity is the presence of the 4ʹ methyl group in the butenolide moiety. All known non-synthetic SLs possess a methyl-substituted carbon at the 4ʹ position (Nomura et al., 2024). Synthetic SLs without this methyl group (i.e. desmethyl-SLs) cannot induce D14-mediated inhibition of bud outgrowth in *Pisum sativum* (Boyer et al., 2012), nor D14-mediated suppression of hypocotyl elongation in *A. thaliana* (Yao et al., 2021). D14 does not undergo thermal destabilisation in the presence of desmethyl butenolides such as dGR24, but is highly sensitive to GR24 with the 4ʹ methyl group (Yao et al., 2021). Conversely, desmethyl butenolides are preferred by KAI2, as indicated by thermal destabilisation, ligand binding, and hydrolysis assays (Yao et al., 2021). In physiological assays such as the inhibition of seedling hypocotyl elongation, dGR24 is uniquely bioactive through KAI2 (Yao et al., 2021).

Enigmatically, D14 hydrolyses desmethyl butenolides at a greater rate than methyl-substituted equivalents (Yao et al., 2018a; Yao et al., 2021). However, as desmethyl butenolides are biologically inactive via D14, and D14 protein is only thermally destabilised by 4ʹ-methyl-substituted butenolides (Yao et al., 2021), desmethyl butenolides presumably behave as simple enzymatic substrates that are turned over rapidly, but cannot activate D14 to initiate signalling.

The divergence in substrate preference between D14 and KAI2 raises the question of how and when these differences arose during the evolution of land plants. Given that D14 is predicted to be a relatively recent duplicate of KAI2, one might hypothesise that the earliest D14 proteins would have had a substrate preference more like KAI2. Ancestral sequence reconstruction (ASR) is a molecular evolution technique for studying the evolutionary trajectory of proteins. The technique relies on multiple sequence alignment and phylogenetic tree construction to generate a sequence that may have existed in the common ancestor (Thornton, 2004). This method serves as an alternative to comparing many orthologous proteins; instead of characterising individual extant orthologues, one could instead study a representative ancestral sequence to infer the ancestral function of the group as a whole.

One of the primary uses of ASR is to gain insight into evolution at a molecular level. For example, ASR has been used *in planta* to understand the evolution of self-incompatibility mechanisms in Brassicaceae through the generation of an ancestral S-Locus Receptor Kinase, showing that the divergence of two allelic variants was asymmetrical: one variant resembles the specificity of the ancestor, while the other is neofunctionalised with novel ligand specificity (Chantreau et al., 2019). Another study has used ASR to characterise the evolution of binding pockets in α/β-hydrolases, using soluble epoxide hydrolases from plants as a representative group (Bzówka et al., 2022). In addition to studying molecular evolution, the technique has been used for protein engineering, as a means of exploiting the features associated with ASR proteins. ASR typically produces proteins that have increased thermostability (Gaucher et al., 2003; Gaucher et al., 2008; Wijma et al., 2013; Gumulya et al., 2018), allowing for modified versions of the protein to tolerate mutations that are destabilising but functionally useful (Risso et al., 2014). In addition to increased thermostability, or perhaps even as a direct result of it, ASR-designed proteins tend to display elevated recombinant expression levels, making it easier to scale protein production for industrial settings (Lei et al., 2023), or at least for ease of experimentation in a smaller-scale laboratory setting. Another major desirable trait for protein engineering is the activity of an enzyme, both with respect to its turnover rate (Gumulya et al., 2018) and its specificity (Siddiq et al., 2017). ASR has again proved to be useful in heightening enzyme activity against a broader range of substrates, leading to enhanced outcomes for enzyme-dependent industrial processes and biopharmaceutical development (Risso et al., 2013; Barruetabeña et al., 2019).

In this study, we describe an ancestral reconstruction of D14 from seed plants. We found that the ancestral protein was structurally highly similar to extant D14 proteins, exhibited traits desirable for an engineered protein, and behaved as a bona fide strigolactone receptor *in planta*. However, we did not find any evidence for wider substrate specificity, relative to D14 from *Arabidopsis thaliana*.

## MATERIALS AND METHODS

### Generation of ancestral sequences

The ancestral D14 sequence was derived manually based on sequence alignment. For 221 out of 264 positions, there was clear conservation of amino acid usage within the DLK4+Eu-D14 clade, and thus a straightforward decision for those positions in the ancestral sequence. For the remaining 43 positions, there was no clear consensus within the DLK4+EuD14 clade, so we instead considered consensus in larger number of groups of proteins; Eudicot D14, Monocot D14, Basal angiosperm D14, Gymnosperm D14, gymnosperm DLK4, gymnosperm DLK4A, gymnosperm DLK4B, fern DDK and lycophyte DDK. Based on the distribution of residues across these groups, it was usually possible to establish a likely ancestral residue. For instance, for position 43, angiosperm D14 proteins have a consensus for tyrosine (Y), but both gymnosperm D14 and gymnosperm DLK4 proteins have a consensus for phenylalanine (F). Thus angiosperm 43Y is derived, and 43F is likely ancestral. For position 128, DLK4 sequences have a consensus for glycine (G) while EuD14 has a consensus for aspartic acid (D); however, fern DDK proteins (the sister group to DLK4+EuD14) has a consensus for aspartic acid, implying that the ancestral residue is 128D. In this way, we could assign a plausible ancestral residue for a further 39 positions. For the remaining 4 positions, there was no clear pattern of residues across the clades, and we selected the most frequent amino acid as the consensus across those groups.

### Molecular cloning

Coding sequences for ancD14 and AtD14 were codon optimised for *E. coli* (GenScript GenSmart) and ordered as synthetic double-stranded DNA fragments (Integrated DNA Technologies) with overhangs homologous to the pSUMO-Amp expression vector. For recombinant expression, these fragments were assembled into the linearised pSUMO backbone via Hot Fusion assembly (Fu et al., 2014). Clones were screened via colony PCR, restriction digest, and Sanger sequencing.

To generate plant binary vectors, coding sequences were extracted from pSUMO-Amp expression vectors via proof-reading PCR, with amplicons subsequently assembled into the pMDC43 backbone (Curtis and Grossniklaus, 2003), also via Hot Fusion assembly. Constructs were verified via colony PCR, restriction digest, and Sanger sequencing.

### Recombinant expression and purification

Expression vectors were transformed into chemically competent Rosetta *E. coli* (Novagen) and plated onto LB agar plates containing carbenicillin (100 μg/mL) and chloramphenicol (34 μg/mL). Three colonies from each plate were used to inoculate 5 mL LB containing the same antibiotics, grown at 30 °C/180 rpm overnight as a starter culture, of which 450 μL was used to inoculate 450 mL LB containing carbenicillin (100 μg/mL). Cultures were grown at 30 °C/180 rpm until an OD_600_ >0.5, chilled to 16 °C, and induced with 0.1 mM isopropyl β-D-1 thiogalactopyranoside (IPTG). Cultures were further incubated at 16 °C/180 rpm for a further 16 h and cells collected the following morning via centrifugation (5,000× *g*, 15 minutes). Cell pellets were stored at −70 °C.

Proteins were purified at 4 °C via affinity chromatography with 20 mL gravity columns. Cells were lysed by resuspending pellets in 12.5 mL lysis buffer (20 mM HEPES, 150 mM NaCl, 10% (v/v) glycerol, 20 mM imidazole, 1× BugBuster (Merck), pH 7.5). Lysates were complexed with 2 mL (settled volume) Ni-NTA resin (BioRad) for the purpose of protein assays, or TALON cobalt resin (TaKaRa) for the purpose of crystallisation, for 1 hour with gentle rotation. Columns were washed with 20 mL wash buffer (20 mM HEPES pH 7.5, 150 mM NaCl, 10% (v/v) glycerol, 20 mM imidazole, pH 7.5) twice. Proteins were eluted in 10 mL elution buffer (20 mM HEPES, 150 mM NaCl, 10% glycerol (v/v), 200 mM imidazole, pH 7.5). Imidazole was diluted at least 1000-fold via buffer exchange (with 20 mM HEPES, 150 mM NaCl, 10% (v/v) glycerol, pH 7.5) in 10 kDa centrifugal filter units (Amicon Ultra-15 10,000 NMWL). For crystallisation purposes, proteins were dialysed overnight in the presence of SUMO protease UlpIto remove the 6xHIS-SUMO leader sequence, and purified by passing through the cobalt affinity resin again and collecting flow-through only. Proteins were then concentrated by ultrafiltration using centrifugal filter units. Purity was assessed by SDS-PAGE, and protein concentration was estimated via NanoDrop, measuring OD_280_, or by Bradford assay (Bio-Rad) to directly compare protein yields (Figure S4).

### Protein crystallisation

AncD14 crystals were grown by vapour-diffusion in sitting drop format. AncD14 protein sample at 10 mg/mL was mixed with reservoir solution in a 1:1 ratio to produce a total drop volume of 0.4 μL and equilibrated at 293 K over a reservoir containing 60 μL of 25% (w/v) PEG 1500. Small square bipyramidal crystals grew after ∼ 3-4 weeks to a maximum size of ∼ 100 μm (Figure S5).

### X-ray diffraction data collection

Single crystals were cryoprotected by transferring into Paratone-N (Hampton Research) before being frozen in liquid nitrogen. Diffraction data were collected at the Australian Synchrotron MX1 beamline (Cowieson et al., 2015) at 100 K using an X-ray energy of 13 keV. Data on a single crystal were collected continuously for 36 seconds using an EIGER2 X 9M direct single-photon counting detector (Dectris) while rotating the crystal through 360° such that the final dataset contained 3600 frames, each containing data from rotation through a φ angle of 0.1°.

### Data processing and refinement

Data were indexed using *XDS* (Kabsch, 2010), and scaled and merged in spacegroup *P* 4_3_2_1_2 using *AIMLESS* (Evans and Murshudov, 2013). The structure was solved by molecular replacement using *phaserMR* (McCoy et al., 2007) with the X-ray crystal structure of *A. thaliana* KAI2 (PDB ID: 4HTA) (Bythell-Douglas et al., 2013) as the search model. The model was initially rebuilt in *Coot* (Emsley et al., 2010) to correct for sequence differences between KAI2 and ancD14 before subsequent rounds of automated and manual refinement using *phenix.refine* (Afonine et al., 2012) and *Coot*, respectively, using data to a resolution of 1.95 Å. Refinement of anisotropic B-factors was restricted to protein atoms contained in three translation-libration-screw (TLS) groups defined automatically using *phenix.refine*. Data collection and refinement statistics are listed in Table 1. Atomic coordinates and structure factors are deposited online at the RCSB Protein Data Bank under accession number 7UKB.

**Table 1:** X-ray data collection and refinement statistics.

|  |  |
| --- | --- |
| **Data Collection:**<sup>a</sup> |  |
| Space group | *P*4<sub>3</sub>2<sub>1</sub>2 |
| *a*, *b*, *c* (Å) | 58.79, 58.79, 146.07 |
| $\alpha$, $\beta$, $\gamma$ (°) | 90, 90, 90 |
| Resolution range (Å) | 48.69–1.95 (1.99–1.95)<sup>b</sup> |
| Total No. of reflections | 511,574 (34,144) |
| No. of unique reflections | 19,665 (1333) |
| Completeness (%) | 99.9 (98.0) |
| Redundancy | 26.0 (25.6) |
| $\langle I/\sigma(I) \rangle$ | 11.0 (1.0) |
| *R*<sub>p.i.m.</sub> | 0.047 (0.617) |
| *CC*<sub>1/2</sub> | 0.999 (0.595) |
| Wilson *B*-factor (Å<sup>2</sup>) | 25.5 |
| **Refinement:** |  |
| Resolution range (Å) | 45.797–1.946 (2.016–1.946)<sup>b</sup> |
| Completeness (%) | 99.8 (98.2) |
| No. of reflections, working set | 18578 (1794) |
| No. of reflections, test set | 989 (88) |
| Final *R*<sub>work</sub> | 0.1842 (0.2691) |
| Final *R*<sub>free</sub> | 0.2325 (0.3255) |
| *No. of non-H atoms:* |  |
| Protein | 2055 |
| Ion | 1 |
| Water | 237 |
| Total | 2293 |
| *R.M.S. deviations:* |  |
| Bonds (Å) | 0.0016 |
| Angles (°) | 0.46 |
| *Average B-factors (Å<sup>2</sup>):* |  |
| Protein | 30.54 |
| Ion | 29.28 |
| Water | 36.69 |
| *Ramachandran plot:* |  |
| Most favoured (%) | 97.32 |
| Allowed (%) | 2.68 |
| Outliers (%) | 0.00 |

<sup>a</sup>Data were collected from a single crystal. <sup>b</sup>Values for the outer shell are given in parentheses.

### Structure Analysis

Geometric measurements and molecular graphics were generated using PyMOL (Schrodinger, 2015) and UCSF ChimeraX (Meng et al., 2023). Root-mean-square deviations calculated with the “align” function of PyMol used all atoms. Dihedral (χ) angles were measured using “get_dihedral” in PyMOL. Distances between the catalytic Ser and His residues (at the OG and NE2 atoms, respectively) were measured using the “Distance” function of ChimeraX. Pocket volumes were estimated using CASTpFold (Ye et al., 2024).

### Chemical synthesis

Synthesis of GR24 and dGR24, and separation of their constituent enantiomers, were carried out as previously described (Mangnus et al., 1992; Scaffidi et al., 2014; Yao et al., 2021). Synthesis of YLG (Tsuchiya et al., 2015) and dYLG (Yao et al., 2018a) were also as described previously.

### Hydrolysis assays

Hydrolysis assays were performed in technical triplicates using 96-well black assay plates (Greiner 655900). Proteins were diluted to 100 μg/mL in hydrolysis buffer (100 mM HEPES, 150 mM NaCl, pH 7). Fluorescent hydrolysis substrates (Yoshimulactone Green (YLG) or desmethyl Yoshimulactone Green (dYLG)) were serially diluted in DMSO to form 1000× stocks, which were diluted in hydrolysis buffer to 1.11× reaction conditions immediately prior to dispensing on the assay plate. Final substrate concentrations ranged from 0 to 20 μM. With the 96-well plate on ice, proteins were dispensed 10 μL (1 μg) per well, followed by 90 μL of the substrate. Immediately, the plate was loaded onto a plate reader at 25 °C (Biotek Synergy HTX), followed by a 20-second orbital shaking step, then measurement of fluorescence (excitation/emission of 485/520 nm) once per minute, for 30 minutes. Blank-subtracted fluorescence values (from wells without protein) were analysed in GraphPad Prism, using nonlinear regression with the “Michaelis-Menten” model. The initial hydrolysis rate was defined as the change in blank-subtracted fluorescence units over the linear phase of the reaction.

### Hydrolysis competition assays

Competition assays were performed as described for hydrolysis assays, but with dYLG only at a fixed concentration of 5 μM. In addition to dYLG, enantiomers of GR24 (GR24^5DS^, GR24*^ent^*^−5DS^) or dGR24 (dGR24^5DS^, dGR24*^ent^*^−5DS^) were included at varying concentrations from 0 to 50 μM to inhibit the hydrolysis of dYLG, thereby decreasing the rate of fluorescence. Data analysis was performed in GraphPad Prism, using nonlinear regression with the “[Inhibitor] vs. response – Variable slope (four parameters)” model.

### Differential scanning fluorometry

DSF assays were performed in technical quadruplicates using sealed 384-well white assay plates (Axygen PCR-384-LC480-W). Each well was prepared with a final reaction condition of 20 μM protein, 5× SyPro Tangerine Dye (ThermoFisher S12010), 20 mM HEPES pH 7.5, 150 mM NaCl, 5% acetone or DMSO (as a solvent for the substrate), and a varying concentration of a GR24 or dGR24 enantiomer (GR24^5DS^, GR24*^ent^*^−5DS^, dGR24^5DS^, or dGR24*^ent^*^−5DS^), by first mixing and dispensing protein, dye, and buffer, and then mixing and dispensing the ligands and buffer. Reactions were incubated in the dark at ambient temperature for 20 min prior to loading on a LightCycler 480 (Roche), with a ramp rate of 0.2 °C/s from 20 °C to 80 °C, and collecting fluorescence with excitation/emission of 483/640 nm. Quadruplicate data were averaged and plotted using GraphPad prism.

### Complementation of *Arabidopsis thaliana d14* mutant

Binary vectors containing either the *ancD14* or *AtD14* coding sequence were transformed into competent GV3101 *Agrobacterium tumefaciens* via freeze-thaw, with positive transformants being verified via colony PCR. Homozygous *kai2-2 d14-1* double mutants in *A. thaliana* were transformed via floral dip. Primary transformants were selected on 0.5 × MS agar supplemented with 20 μg/mL hygromycin B. T_2_ offspring were screened for 3:1 segregation of resistance to hygromycin B, and further propagated to the T_3_ generation, in which homozygotes were selected that demonstrated 100% resistance to hygromycin B. Three independent transformed lines were generated from each construct. Transformed lines were verified at the gene, transcript, and protein level, by PCR genotyping, RT-qPCR, and immunoblotting.

### Immunoblotting

Total protein was extracted from 7-day-old *A. thaliana* seedlings, which were snap-frozen in liquid nitrogen and homogenised with two Zirconox beads in a mixer mill (30/s). Proteins were extracted by applying 150 μL extraction buffer (50 mM Tris pH 7.5, 150 mM NaCl, 10% glycerol, 0.1% Tween-20, 1 mM dithiothreitol, 1 mM phenylmethylsulfonyl fluoride, 1× Complete Protease Inhibitor Cocktail (Roche)) to 100 mg homogenised seedling tissue and removing cell debris via centrifugation at 4 °C (12,000 × *g*, 10 min; followed by 15,000 × *g*, 10 min). Protein concentration was estimated by Nandrop A280, using the BSA standard, and 80 μg protein extract was separated via 12% SDS-PAGE. Proteins were transferred to a polyvinylidene fluoride membrane using a Trans-Blot Turbo (Bio-Rad).

Immunoblots were blocked with 1× TBS-T (20 mM Tris-HCl, 150 mM NaCl, 0.1% Tween-20, pH 7.5) supplemented with 2% BSA at room temperature for 2 hours. Antibodies were prepared by dilution to 10 mL with 1 × TBS-T with 0.2% BSA and incubated with the immunoblot at 4 °C with gentle rocking for at least 2 hours. Immunoblots were washed five times (5 minutes per wash) with 1 × TBS-T between each antibody incubation. To quantify transgene expression, GFP was probed using rabbit anti-GFP (Invitrogen A11122), followed by goat anti-rabbit IgG HRP (ThermoFisher 31460), and imaged using Clarity Western ECL substrate (Bio-Rad). The immunoblot was then re-blocked as above for a further 2 hours and probed for actin mouse anti-actin (Sigma-Aldrich A0480) followed by goat anti-mouse IgG AP (Pierce 31321), imaged using ImmunStar AP substrate (Bio-Rad). Chemiluminescence was imaged in both instances using an ImageQuantRT ECL system (GE Healthcare).

### Transcript analysis

Total RNA was extracted from 4-day-old *A. thaliana* seedlings using the Spectrum Plant Total RNA Kit (Sigma-Aldrich), following ‘Protocol A’ as described by the manufacturer. RNA was then treated with TURBO DNase (Life Technologies) and used in a cDNA synthesis reaction using the LunaaScript RT SuperMix Kit (New England Biolabs). RT-qPCR was performed using threefold-diluted cDNA with LUNA Universal qPCR Master Mix (New England Biolabs).

### Hypocotyl elongation assays

Surface-sterilised seeds were sown onto 0.5 × MS agar plates (pH = 5.9) containing GR24 molecules diluted from a 1000× stock in DMSO, and stratified in darkness at 4 °C for 72 hours. Mock-treated plates included an identical volume of DMSO. Plates were then moved to 21 °C and exposed to white light for 3 hours, followed by a further 21 hours of darkness, and then 4 days of continuous red light provided by light emitting diodes (λ_max_ 660 nm; 10 μmol m^−2^ s^−1^). Seedlings were randomly selected onto a new agar plate, arranged such that the hypocotyls remained flat on the surface. Hypocotyl lengths were measured using ImageJ (https://imagej.net/ij/).

### Shoot branching & plant height assays

Seedlings were germinated from surface-sterilised seed on 0.5 × MS agar, stratified in darkness at 4 °C for 72 hours, then grown under constant white light for one week at 21 °C. For mature growth, 11 seedlings of each genotype were moved to a mixture of coconut coir, perlite, and vermiculate (6:1:1), grown under long-day conditions. Shoot branching, plant height, and flowering time were recorded daily.

## RESULTS

### Inference of an ancestral D14 sequence

To generate a plausible ancestral D14 (ancD14) sequence, a large dataset of DDK (D14/DLK2/KAI2) protein sequences was considered (Bythell-Douglas et al, 2017). Sequences were compared within the DLK4 + euD14 clade identified in Bythell-Douglas et al (2017), which contains all true D14 sequences in seed plants (as opposed to DLK2 and DLK3 sequences, which are more divergent and functionally distinct). Within this group, we looked at each core amino acid position identified in Bythell-Douglas et al (2017), and assessed whether the residue present at that position was highly conserved (found in 75% of sequences in the group), conserved (found in 50% of sequences in the group), or showed minority consensus (found in 30% of sequences in the group, and the most common residue at that position). We then used these to determine the most plausible residue present at each position in the ancestral sequence of the DLK4 + eu-D14 clade, at the base of seed plants. We settled on a single ancestral sequence that represents a plausible consensus of seed plant D14 proteins (Figure S1; Supplemental Data 1).

### Ancestral D14 complements the *d14* mutation of *Arabidopsis thaliana*

To test if ancD14 can fulfill the role of its counterpart in *Arabidopsis thaliana*, the ancD14 coding sequence was cloned into the plant binary vector pMDC43 (Curtis and Grossniklaus, 2003) to generate a protein fusion with GFP at the N-terminus. Two independent transgenic lines homozygous for the *35S:GFP-ancD14* transgene were generated in the *kai2-2 d14-1* double mutant background, alongside an equivalent line expressing GFP-AtD14 as a control. Expression in the double mutant and under a constitutive promoter allowed for an assessment of functional orthogonality, in case ancD14 should have any unexpected KAI2-related activity. The functionalities of ancD14 and AtD14 were compared *in planta* with respect to plant height and shoot branching phenotypes.

The *kai2-2 d14-1* double mutant exhibited a dwarf stature compared to wild-type L*er* throughout the reproductive phase of growth (Figure S2A). The *35S:GFP-AtD14* transgene largely recovered this dwarf phenotype, although there was a notable delay in flowering that presumably results from the *kai2-2* mutation, because this phenotype was not affected by either of the transgenes (Figure S2C). Accordingly, plant height was evaluated 14 days after flowering initiated, rather than plant age (Figure S2B). When compared to *35S:GFP-AtD14*, one of the two *35S:GFP-ancD14* lines was slightly shorter on average, but both were significantly taller than *kai2-2 d14-1* (Figure 1A). Therefore, with respect to plant height, we conclude that ancD14 offers near-complete complementation of the *d14* phenotype in *A. thaliana*.

**Figure 1:**
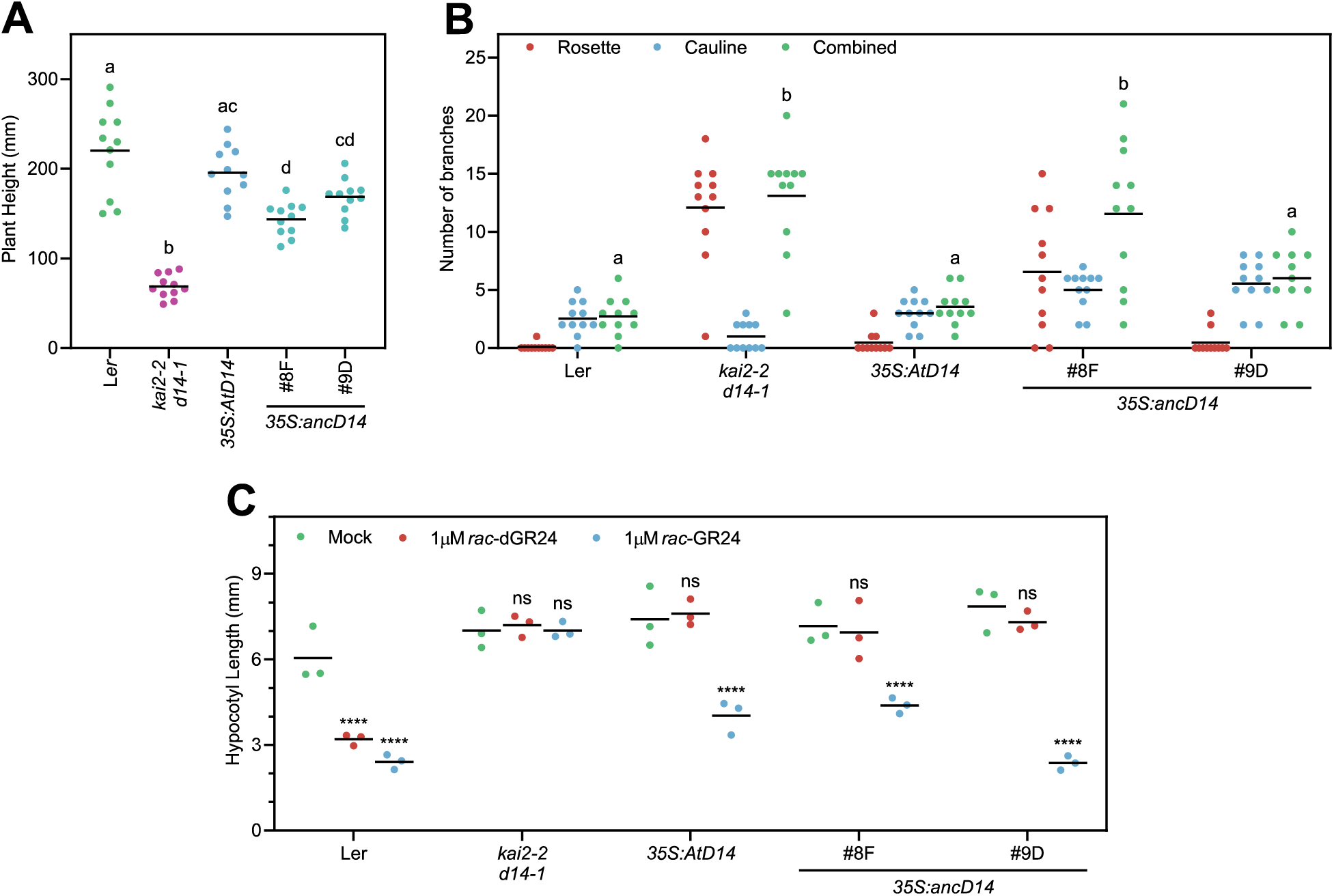
Phenotypes of *Arabidopsis thaliana* constitutively expressing ancD14. (**A**) Scatter plot of plant height evaluated 18 days after the emergence of the primary inflorescence. Significantly different groups denoted by lowercase letters; *n* = 11. (**B**) Scatter plot of shoot branching observed in 80-day-old plants. Significantly different groups denoted by lowercase letters for the combined number of branches only; *n* = 11. (**C**) Suppression of hypocotyl elongation in relevant transgenic lines by synthetic strigolactone analogues. Each data point represents the mean of *n* = 3 independent replicate experiments, each comprising ≥ 20 seedlings per treatment. Results from individual replicate experiments are shown in Figure S2D; ****, p < 0.0001; ns, not significant (two-way ANOVA with multiple pairwise comparisons for simple effects within rows)

The *kai2-2 d14-1* double mutant displays increased shoot branching relative to L*er*, and *35S:GFP-AtD14* was able to revert this mutant phenotype as expected (Figure 1B). The two lines expressing GFP-ancD14 exhibited different shoot branching phenotypes, with line #9D also restoring branching back to that of L*er*, but individuals of line #8F had very variable branching phenotypes, such that overall #8F was not significantly different from *kai2-2 d14-1* in this respect. From these data, we conclude that ancD14 can partially to fully recapitulate the *d14* phenotype in *A. thaliana*.

The transgenic Arabidopsis lines expressing GFP-AtD14 and GFP-ancD14 were also assayed for their ability to respond towards different ligands, by measuring the suppression of hypocotyl elongation (Figure 1C). Re-introduction of *AtD14* in the *kai2-2 d14-1* double mutant background allowed for responses to *rac*-GR24, but not to *rac*-dGR24. Both lines expressing ancD14 demonstrated the same pattern, with line #9D showing the greater response commensurate with its stronger expression of the transgene. These data confirm that ancD14 can function as a receptor for SLs in a similar manner to AtD14. No evidence was found that ancD14 could compensate for loss of *KAI2*, based on the elongated hypocotyl phenotype, lack of response to *rac*-dGR24, and the impaired transcription of the downstream marker *DLK2* (Figure S2E).

We suspected that partial complementation and differing phenotypes between the two independent lines expressing ancD14 could reflect differences in transgene expression level. RT-qPCR revealed that line #8F, which exhibited weaker complementation of the observed phenotypes, had lower and less consistent transgene expression at the transcript level compared to the other transgenic lines (Figure S3A). These differences also resulted in weaker expression on the protein level (Figure S3B). These findings provide a plausible explanation for the incomplete and variable phenotypic complementation observed in line #8F.

### Ancestral D14 can bind and hydrolyse butenolides *in vitro*

To understand if ancD14 can act as a receptor and an enzyme towards SLs, recombinant ancD14 was expressed as a SUMO-fusion protein and compared with AtD14 in several protein-based assays. During these experiments, we found that the average yield of SUMO-ancD14 was approximately 5.1 mg/L of bacterial culture, more than double that of SUMO-AtD14 (Figure S4).

D14 and its homologue KAI2 both show hydrolytic activity towards the profluorescent compound Yoshimulactone Green (YLG), with strongly enhanced activities towards desmethyl Yoshimulactone green (dYLG) (Tsuchiya et al., 2015; Yao et al., 2018a; Yao et al., 2021). We compared the activity of ancD14 with AtD14 to identify any changes in enzymatic activity or substrate promiscuity. We observed that ancD14 shows strong hydrolytic activity towards dYLG over YLG, in a similar fashion to AtD14 (Figure 2A). Notably, with concentrations of enzyme kept constant, ancD14 exhibited approximately double the maximal rate of enzymatic activity than AtD14 towards both substrates. However, K_m_ values differed considerably. Although the affinity of ancD14 for dYLG was just two-fold lower than that of AtD14, it was some two orders of magnitude lower for YLG (Figure 2A). Thus, ancD14 is a more active enzyme than AtD14, but displayed much reduced affinity for the YLG substrate that most closely mimics natural strigolactones.

**Figure 2:**
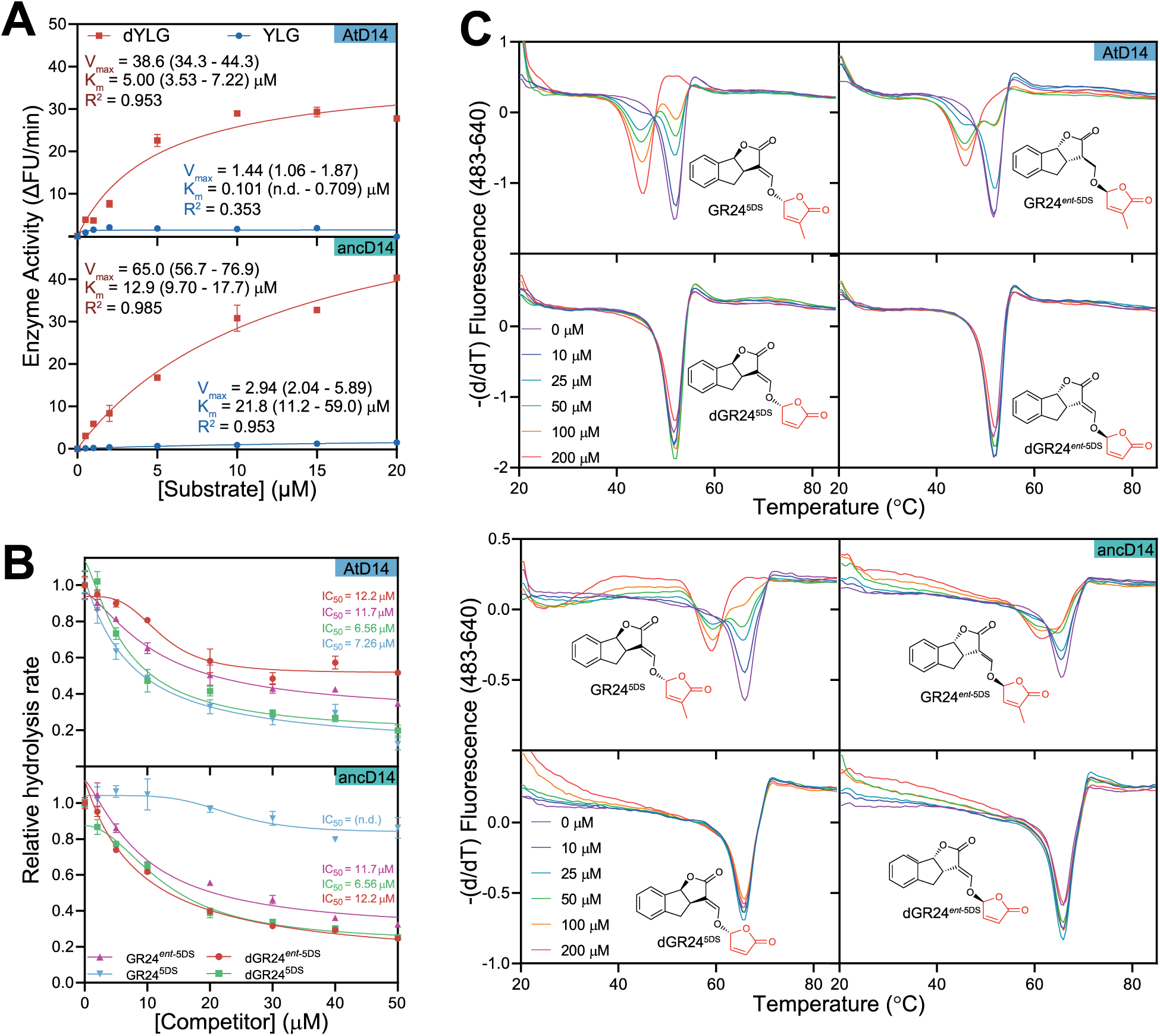
Ancestral D14 interacts with butenolides. (**A**) Hydrolysis of Yoshimulactone Green (YLG) and desmethyl Yoshimulactone Green (dYLG) by equivalent concentrations of AtD14 (top) and ancD14 (bottom). Data points are presented as means ± SE for *n* = 3 technical replicates; Michaelis-Menten values are provided with 95% CI estimates in brackets; n.d., not determinable. (**B**) Suppression of dYLG hydrolysis by GR24 and dGR24 compounds. Hydrolysis of 5 μM dYLG was competed with 0 – 50 μM GR24 or dGR24 enantiomers in the presence of 100 ng protein. Data points are presented as means ± SE for *n* = 3 technical replicates; IC_50_ values are provided with 95% CI estimates in brackets; n.d., not determinable. (**C**) Differential scanning fluorometry of AtD14 (top) and ancD14 (bottom) incubated with 0 – 200 μM of individual enantiomers of GR24 and desmethyl-GR24 (dGR24). Data points are means of *n* = 4 technical replicates; insets show ligand structure.

To investigate the substrate specificity of ancD14 further, we conducted hydrolysis competition assays with constant concentration of dYLG, titrated against increasing concentrations of the constituent enantiomers of GR24 and dGR24. Inhibition of dYLG hydrolysis was used to infer a relative affinity of the enzyme for each enantiomer (Figure 2B). AtD14 activity was inhibited by all four enantiomers, but GR24^5DS^ and dGR24^5DS^ with SL-type 2ʹ*R*-configured D-rings were the most effective, as reported previously (Yao 2021). In contrast, GR24^5DS^ was the only compound that failed to effectively compete with dYLG for hydrolysis by ancD14, suggesting that relative to AtD14, ancD14 does not have a strong affinity for compounds with SL-type stereochemistry.

To obtain more direct evidence of GR24 interacting with ancD14, differential scanning fluorometry was used to screen for thermal shifts indicative of ligand-receptor interactions. AtD14 was thermally destabilised by both GR24^5DS^ and GR24*^ent^*^−5DS^ (but not the dGR24 equivalents) as reported previously (Waters et al., 2015; Yao et al., 2021). Consistent with hypocotyl elongation responses, the same preference was held by ancD14, albeit with a more muted response to GR24*^ent^*^−5DS^ (Figure 2C). In the absence of a ligand, ancD14 displayed a substantially elevated melting temperature (65.9 °C) compared to that of AtD14 (51.7 °C). Nevertheless, GR24^5DS^ lowered the melting temperature of AtD14 by 6.1 °C (from 51.7 °C to 45.6 °C), and of ancD14 by 6.6 °C (from 65.9 °C to 59.3 °C). Despite the large absolute differences in melting temperature, the comparable ligand-induced shifts in melting temperature are consistent with a similar conformational change occurring in both proteins. Furthermore, these data indicate that, like AtD14, ancD14 retains a requirement for a ligand with a methyl-substituted butenolide moiety to trigger this conformational change. Notably, the ligand-induced conformational change in the isolated protein mirrors the ligand bioactivity *in planta*.

### The structure of ancestral D14

To investigate the structural characteristics of the ancestral D14 protein, the crystal structure of the recombinant protein was elucidated to a resolution of 1.95 Å (Figure 3, Table 1). As expected, the structure of ancD14 bears close similarity to other D14 family proteins, and the α/β hydrolase family overall, based on a seven-stranded mostly parallel β-sheet flanked by α-helices, with the active site located between the C-terminal edge of the β-sheet and a largely helical lid domain. All atom root-mean-square deviations range from 0.4 Å for Petunia DAD2 through D14 and KAI2 variants to 1.0 Å for *Bacillus subtilis* RsbQ, in line with sequence similarity (Figure 3B, 3C, 3G, Table S1). The 1.95 Å resolution electron density of ancD14 is generally excellent, revealing clear conformations of side-chains and well-defined solvent molecules. Analysis of the active site geometry reveals features that place ancD14 apart from other D14 structures. For example, the distance between key atoms in the catalytic serine and histidine residues is substantially shorter than other D14 structures (Figure 3E), while the sidechain dihedral angle of serine 95 (χ_1_) which determines the position of the hydroxyl group, is at the extreme of the distribution of χ_1_ angles in the whole family (Figure 3F), with other D14 and DAD2 proteins lying at the other extreme. However, the overall pocket volume was estimated to be typical of that of a eu-D14 protein (Figure 3G). To explore the significance of these observations, close inspection of all available structures of homologues revealed that the active site geometry is influenced by the presence of solvent molecules or ions adjacent to the catalytic serine. In the case of ancD14 a chloride ion is observed at this position, while in the cluster of structures with similar χ_1_ angles a water, chloride or magnesium ion is modelled. Thus, it is possible that the specific detail of active site geometry of α/β hydrolases may be influenced by the crystallisation conditions as much as by homology. Nevertheless, the structure of ancD14 is functionally intact, and displays the conformation and relevant interactions expected for a competent enzyme.

**Figure 3:**
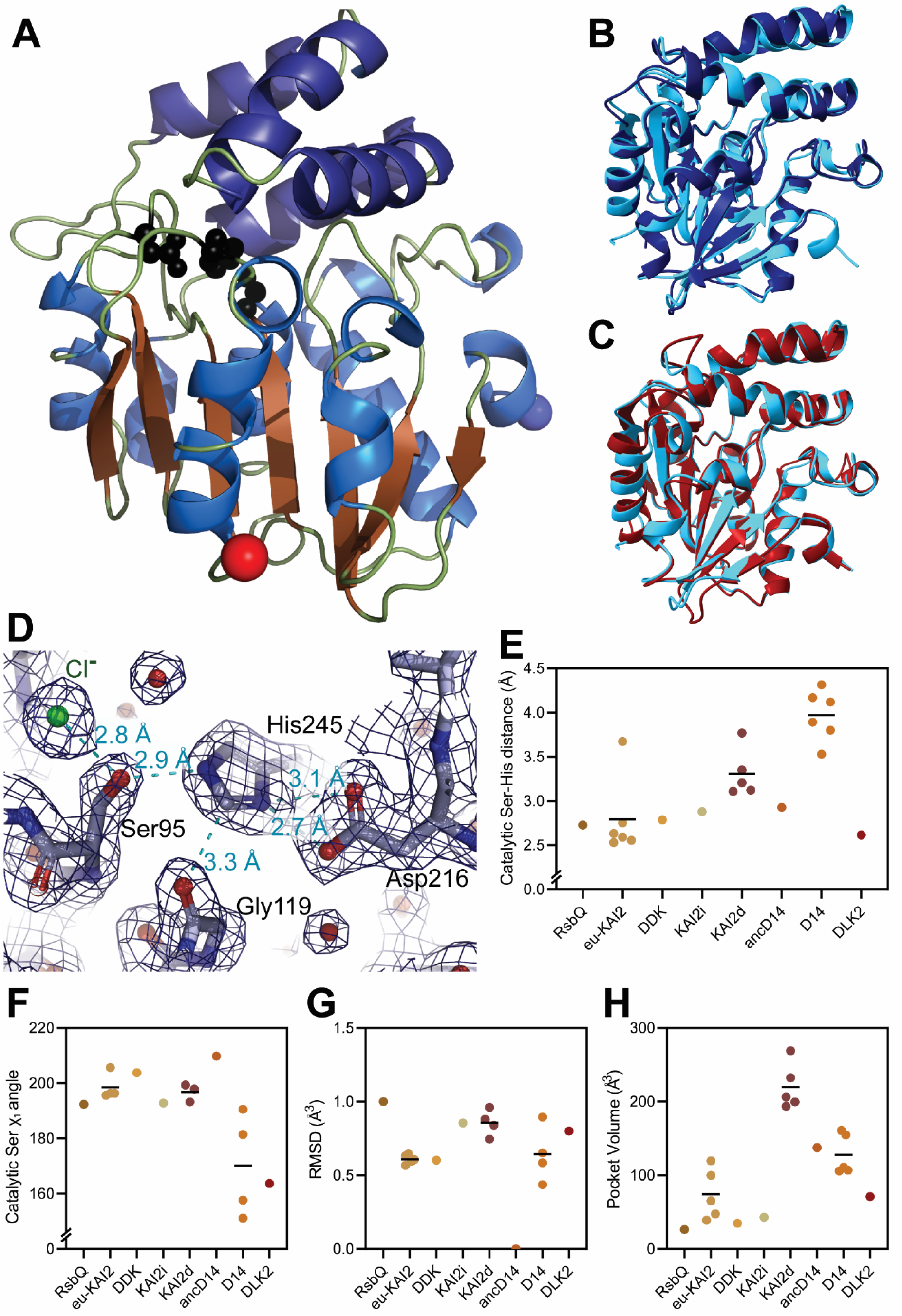
Crystal structure of ancestral D14 (ancD14, PDB 7UKB). (**A**) Cartoon representation of ancD14 coloured by secondary structure (helices, blue; strands, brown). The N- and C-termini are represented by blue and red spheres, respectively. The catalytic triad is represented by black balls-and-sticks. (**B**) Structural alignment of ancD14 and AtD14 (4IH4); RMSD = 0.896 Å. (**C**) Structural alignment of ancD14 and AtKAI2 (4HRX); RMSD = 0.727 Å (**D**) The active site of ancD14, in ball-and-stick representation. Blue mesh, 2mFo-DFc electron density represented at 1.25 σ. Key interatomic distances are indicated by cyan dashed lines. (**E**, **F**, **G**, **H**) Comparison of catalytic serine-histidine distance (E), catalytic serine χ_1_ angle (F), RMSD (compared to ancD14) (G), and pocket volume (H) within KAI2/D14-type homologues. Where several structures are published of one homologue, the mean distance is indicated. Underlying data are provided in Table S1.

## DISCUSSION

### Ancestral D14 substitutes functionally for AtD14

Combined biochemical and biological data indicate that ancD14 can fulfil the same function as AtD14. Accounting for differences in expression level, ancD14 can replicate the biological role of AtD14 in *A. thaliana* with respect to plant height, shoot branching, and GR24-dependent suppression of hypocotyl elongation. The ability of ancD14 to perceive GR24 is further supported at the protein level by DSF, showing a similar thermal destabilisation in response to GR24, but not dGR24, as shown by AtD14. In addition to the responses observed towards GR24 *in planta*, ancD14 was also able to restore phenotypes in *A. thaliana* without any exogenous treatment, indicating that it can perceive natural SLs as well as the synthetic GR24.

Additionally, ancD14 can function not only as a receptor, but as a hydrolase towards the profluorescent butenolides YLG and dYLG. The hydrolysis of dYLG by ancD14 was inhibited by both enantiomers of dGR24 and by GR24*^ent^*^−5DS^, but not by GR24^5DS^, suggesting that this compound is unable to compete with dYLG for the active site. This result is counterintuitive, considering that GR24^5DS^ is thought to resemble the configuration of endogenous SLs most closely, and that GR24^5DS^ is active in DSF. The result is consistent with the very low affinity of ancD14 for YLG relative to AtD14, because both YLG and GR24^5DS^ have a methyl-substituted D-ring. Nevertheless, we have no evidence that desmethyl butenolides are bioactive via ancD14, and therefore the strong rate of dYLG hydrolysis by ancD14 and its apparent affinity for dGR24 do not imply a change in ligand preference towards that of KAI2. Instead, and like AtD14 (Yao 2021), these desmethyl butenolide compounds act only as hydrolysis substrates for ancD14’s enzyme activity, without having the capacity to activate the receptor function by triggering a conformational change. Nevertheless, if the absolute substrate preference has not changed, the reduced affinity of ancD14 towards YLG and GR24^5DS^ suggest that the sensitivity of ancD14 toward endogenous SLs may be weaker relative to AtD14 and other natural receptors, which may be attributed to the atypically compact active site apparent in the structure of ancD14. The reduced effectiveness of ancD14 compared with AtD14 in complementing the *d14* phenotype might result from an impaired affinity for endogenous SLs, but it is also possible that weaker protein-protein interactions with other SL signalling partners such as MAX2 or SMXL/D53 could explain this effect. Screening a greater variety of natural SLs and analogues, both *in vitro* and *in vivo*, would help to assess these uncertainties more comprehensively.

### Ancestral D14 possesses desirable traits for biotechnology

As ancD14 behaves in a comparable manner to AtD14, it serves as a useful tool to exploit its advantageous traits, such as heightened thermostability, elevated expression yield in *E. coli*, and increased enzymatic activity, while preserving the essential functions of a D14 protein. The enhanced expression yield of ancD14 represents an opportunity to scale up D14 protein expression and facilitating the use of D14 in high-throughput mutant screens. The drastically enhanced thermostability also renders ancD14 an ideal platform for introducing further destabilising mutations that might not otherwise be tolerated. These beneficial attributes broaden the utility of the protein in applications such as a biosensor, or as a functional standard with improved useability and storage characteristics.

We did not test the activity of ancD14 in plants beyond *A. thaliana*, so it is not known for certain whether ancD14 can mediate cereal traits such as tillering. However, D14 is highly conserved and considered to be modular within angiosperms; for example, D14 from *Oryza sativa* can fully complement the mutant phenotype when introduced to *A. thaliana* (Yao et al., 2018b). Recently, it was discovered that SL signalling in rice is enhanced by the N-terminal domain (NTD) of D14. Under conditions of low nitrogen availability, phosphorylation of serine residues in the NTD increases the protein’s stability and prolongs signalling lifetime (Hu et al., 2024). However, this NTD is only found in monocot D14 homologues, and likewise the regulatory mechanism that acts via the NTD may be monocot-specific as well. Conceivably, the improved stability of ancD14 could be exploited to allow for a similar trait to be introduced to other crop species. Conversely, the enhanced enzymatic activity of ancD14 could serve to create an SL-decreased environment; for example, by mutating the residues associated with MAX2 interaction to abolish signalling and expressing it under a conditional promoter, SL signalling could be “knocked down” in a context-dependent manner.

### Strigolactone signalling is tolerant of structural variation at the ligand binding site

Overall, the structure of ancD14 is similar to other D14 structures published previously. The most notable point of difference between ancD14 and extant homologues lies in the configuration of the active site. Specifically, ancD14 possesses a compact catalytic triad where the χ_1_ angle of the serine residue and the distance between the serine and histidine residues are more characteristic of a KAI2-type protein. This observation aligns with the evolutionary relationship between the two proteins, but these structural features did not lead to any substantive changes in ligand preference *in vitro* or functional performance *in planta*. This remains consistent with divergent KAI2 proteins from the moss *Physcomitrium patens* or the lycophyte *Selaginella moellendorffii*, which likely serve as SL receptors because non-seed plants lack D14-like sequences or predicted structures (Waters et al., 2015; Bythell-Douglas et al., 2017; Lopez-Obando et al., 2021). It is also noteworthy that the substrate preferences are easily altered: for example, as few as three substitutions can convert a karrikin receptor into a strigolactone receptor (Arellano-Saab et al., 2021). If the ability to perceive SLs does not require a highly specific structural conformation at the active site, perhaps the ability to perceive SLs is instead more dictated by the capacity to recruit signalling partners, specifically MAX2 and/or SMXL proteins.

### Insights and limitations of ASR for understanding D14 evolution

It is challenging to draw definitive conclusions about protein evolution from a single ancestral reconstruction alone. ASR relies heavily on accurate phylogenetic trees and sequence alignments, and different models of evolution can infer different ancestral states. To have confidence in understanding D14 evolution, multiple independent ancestral reconstructions should be studied to mitigate uncertainties associated with any potential phylogenetic inaccuracies and model dependencies.

Nonetheless, our reconstruction provides preliminary evidence that D14-specific traits emerged quickly after the emergence of D14 itself in seed plants. The ligand specificity of ancD14, regarding the presence of the 4ʹ methyl group and 2ʹ stereochemistry of the D-ring, appears to mirror that of modern D14 receptors rather than KAI2 receptors. Furthermore, ancD14 can regulate traits specific to AtD14 *in planta*, but not those mediated by AtKAI2 such as flowering time or the upregulation of *DLK2* transcription (Figure S2C, S2E). These findings collectively suggest that the functional specialisation and ligand specificity of D14 evolved early and have remained relatively stable throughout its evolutionary history, reflecting an early adaptation of D14 to a SL-specific signalling role. The ability of ancD14 to perceive GR24 *in planta* also provides evidence that interactions between D14 and MAX2 or SMXL-family proteins have similarly emerged early and are conserved throughout seed plants.

It has been previously proposed that SL signalling may have more relaxed structural requirements than KAI2-mediated signalling (Bythell-Douglas et al., 2017), in which case attributes such as the more compact active site architecture, or the potential reduced affinity for SLs of ancD14 observed in *in vitro* assays, may reflect a transitional phase in the evolution of SL signalling. Arguably, later evolution might have selected for improved ligand affinity, potentially conferring greater sensitivity and spatial resolution upon SL response.

## SUPPORTING INFORMATION

**Figure S1:**
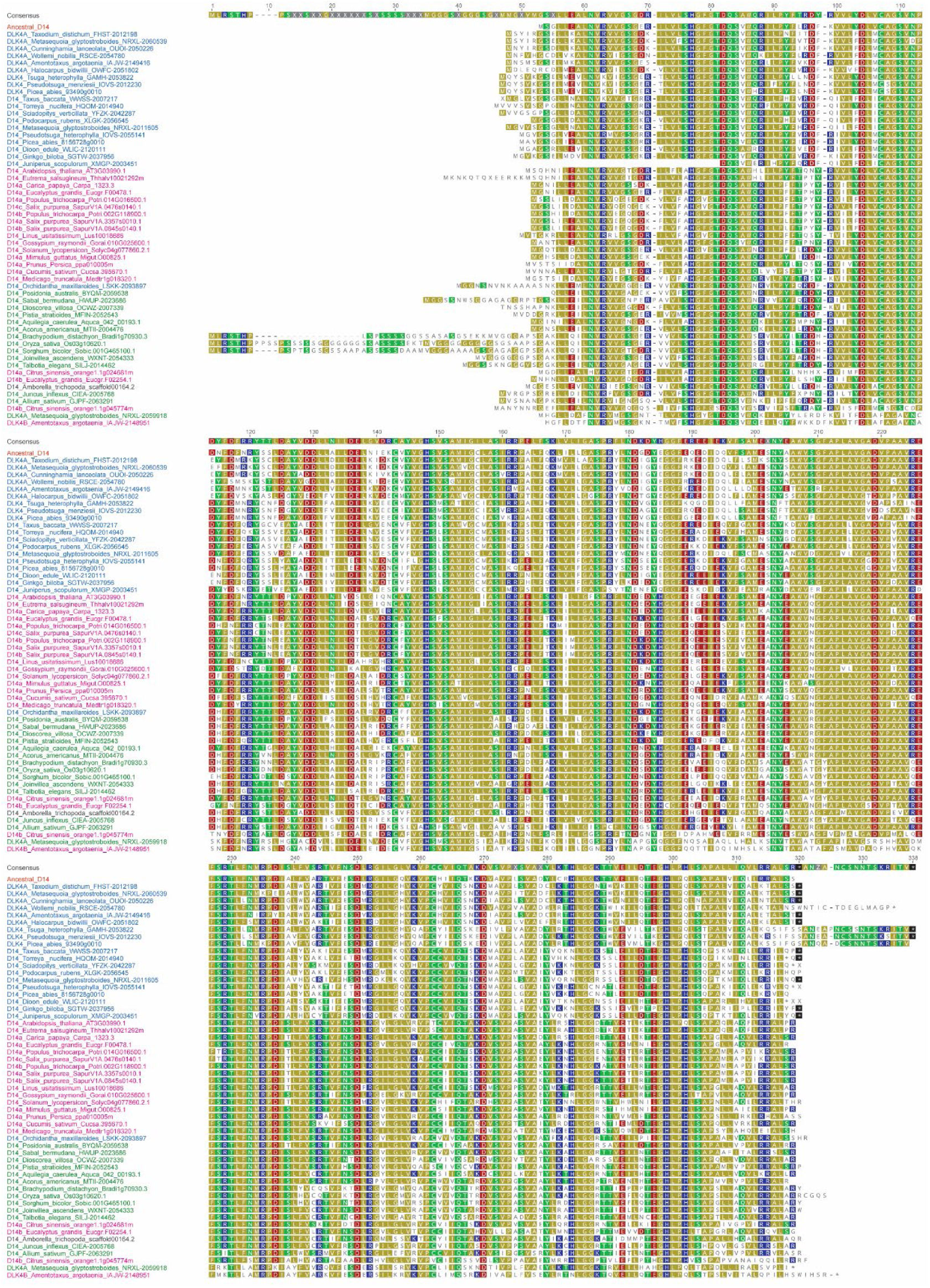
The ancestral-reconstructed D14 sequence aligned to D14 and DLK4 sequences from seed plants. Sequences were aligned by MAFFT v7.49 and are grouped by similarity to each other. Gymnosperms, blue; monocots, green; dicots, fuschia; basal angiosperms, black; ancD14, orange. Amino acid sequences are listed in Supplemental Data 1.

**Figure S2:**
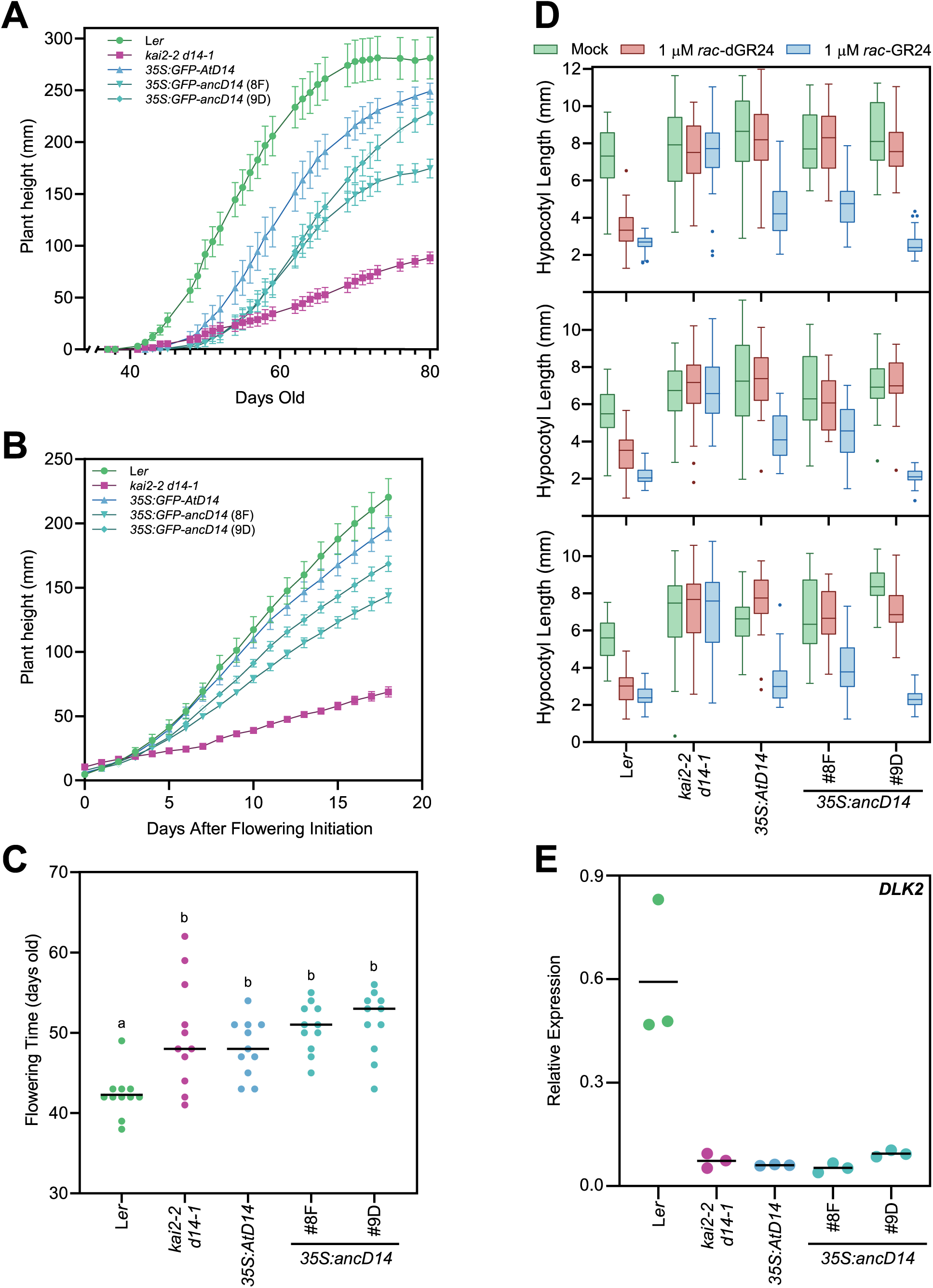
Detailed growth characterisation of Arabidopsis *GFP-ancD14* transgenic lines. Where error bars shown, data are representative of mean ± SE for *n* = 11 biological replicates. (**A**) Plant height over time defined as days post-germination. (**B**) Plant height over time, defined as days post–initiation of flowering. (**C**) Scatter plot of flowering time for each genotype, lines represent the mean for that genotype, lower case letters represent significantly different groups. (**D**) Individual hypocotyl elongation assay replicates. Each panel represents a separate experiment in which *n* ≥ 23 seedlings per genotype-treatment combination were measured. Data presented as a box plot, with outliers (exceeding 1.5 × interquartile range deviation from the median) shown as individual points. (**E**) Relative expression level of *DLK2* transcripts in 9-day-old seedlings quantified by RT-qPCR. Transcript abundance was normalised to the geometric mean of *CACS* and *TIP41L* transcripts; *n* = 3 biological replicates; lines depict means.

**Figure S3:**
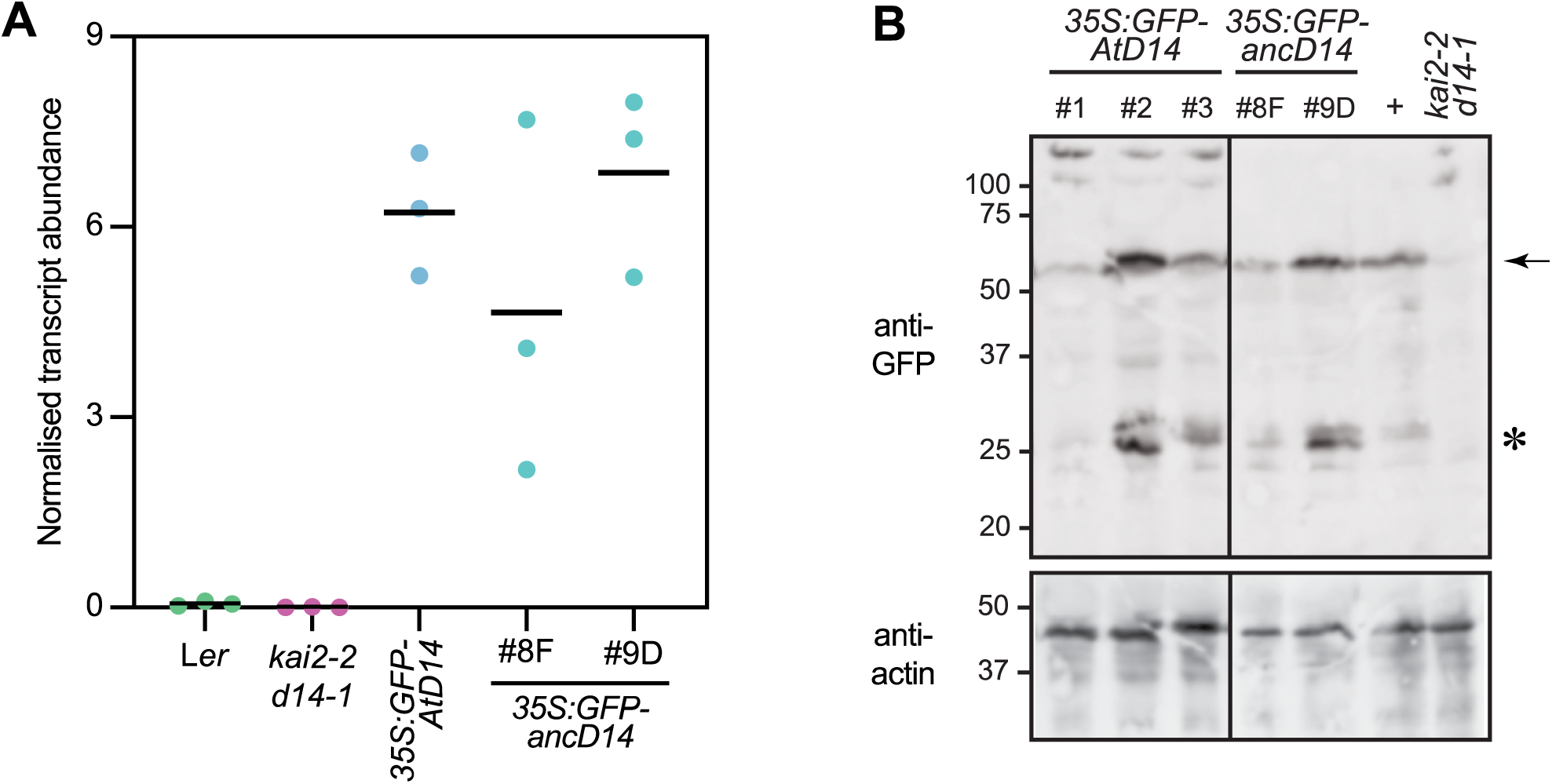
Expression analysis of GFP-D14 proteins in transgenic *Arabidopsis thaliana*. (**A**) Levels of *GFP-AtD14* and *GFP-ancD14* transcripts in stable transgenics, quantified by RT-qPCR using primers anchored in the GFP coding region. Transcript abundance was normalised to the geometric mean of *CACS* and *TIP41L* transcripts. *n* = 3 biological replicates of 9-day-old seedlings; lines depict means. (**B**) Immunoblots targeting the GFP-D14 fusion proteins (upper panel) and actin (lower). Gels were loaded with soluble protein extract from 9-day-old seedlings of two to three independent transgenic lines for each plasmid construct. The blot was probed sequentially with anti-GFP (Invitrogen A11122) and then anti-actin (Sigma A0480) antibodies. The positive control (+) is an extract from Arabidopsis seedlings expressing GFP-BdKAI2 fusion protein (Meng et al., 2021). The arrow depicts the anticipated GFP-D14 fusion protein of ∼60 kDa, and the asterisk indicates an assumed cleavage product of free GFP (∼27 kDa). Line #3 of *35S:GFP-AtD14* was used in this study and in panel A. Vertical line indicates where relevant samples from a larger blot were spliced together for presentation.

**Figure S4:**
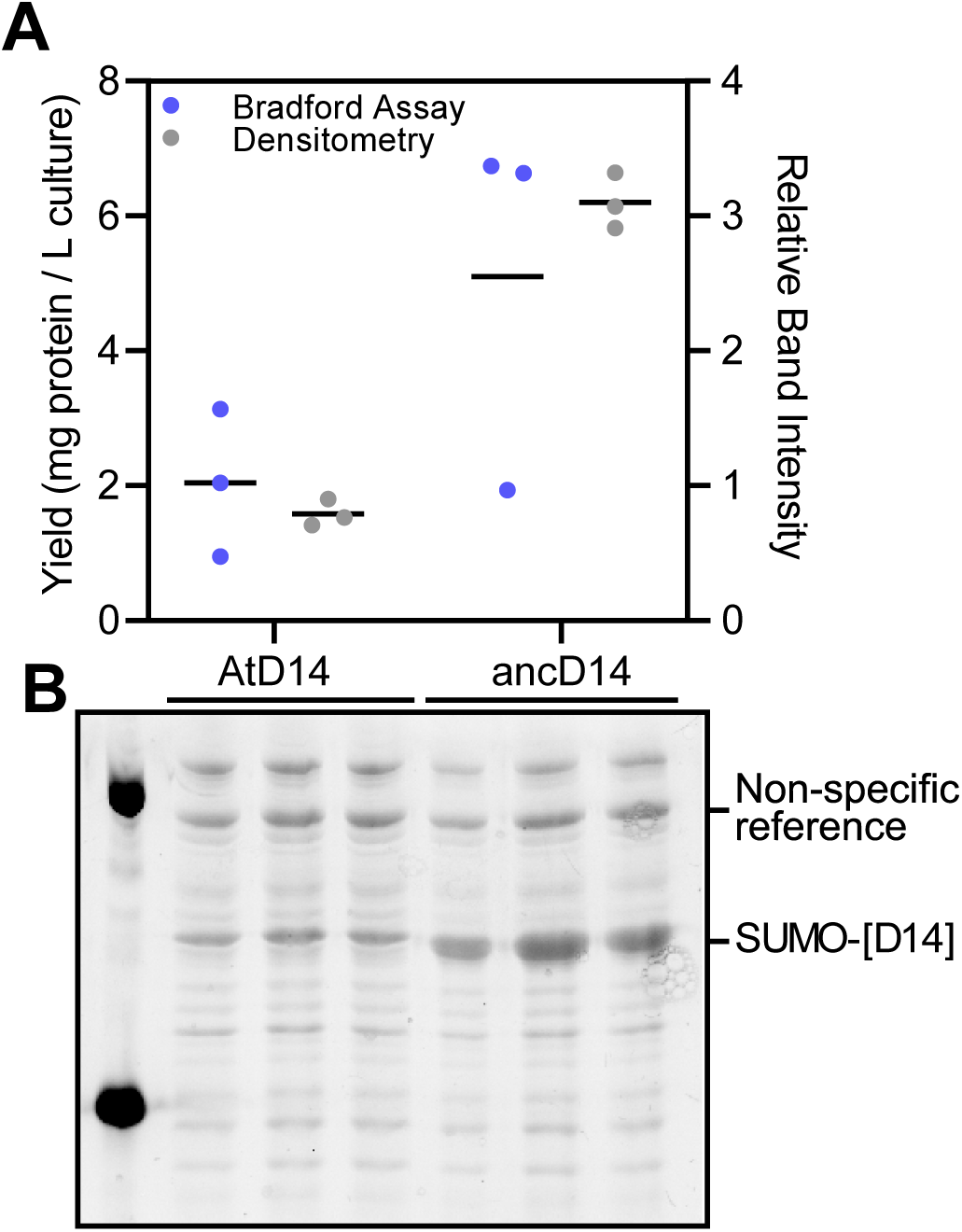
Heterologous expression of AtD14 and ancD14 in *E. coli*. SUMO-fusion proteins were expressed in triplicate 50-mL cultures, and purified from 10 mL samples using a crude magnetic bead chromatography method. (**A**) Estimates of yield were made through Bradford assay interpolated from a standard curve in which known concentrations of SUMO-ancD14 were used (left Y axis), and by densitometry (relative to a non-specific reference band; right Y axis) from the SDS-PAGE gel shown below (**B**).

**Figure S5:**
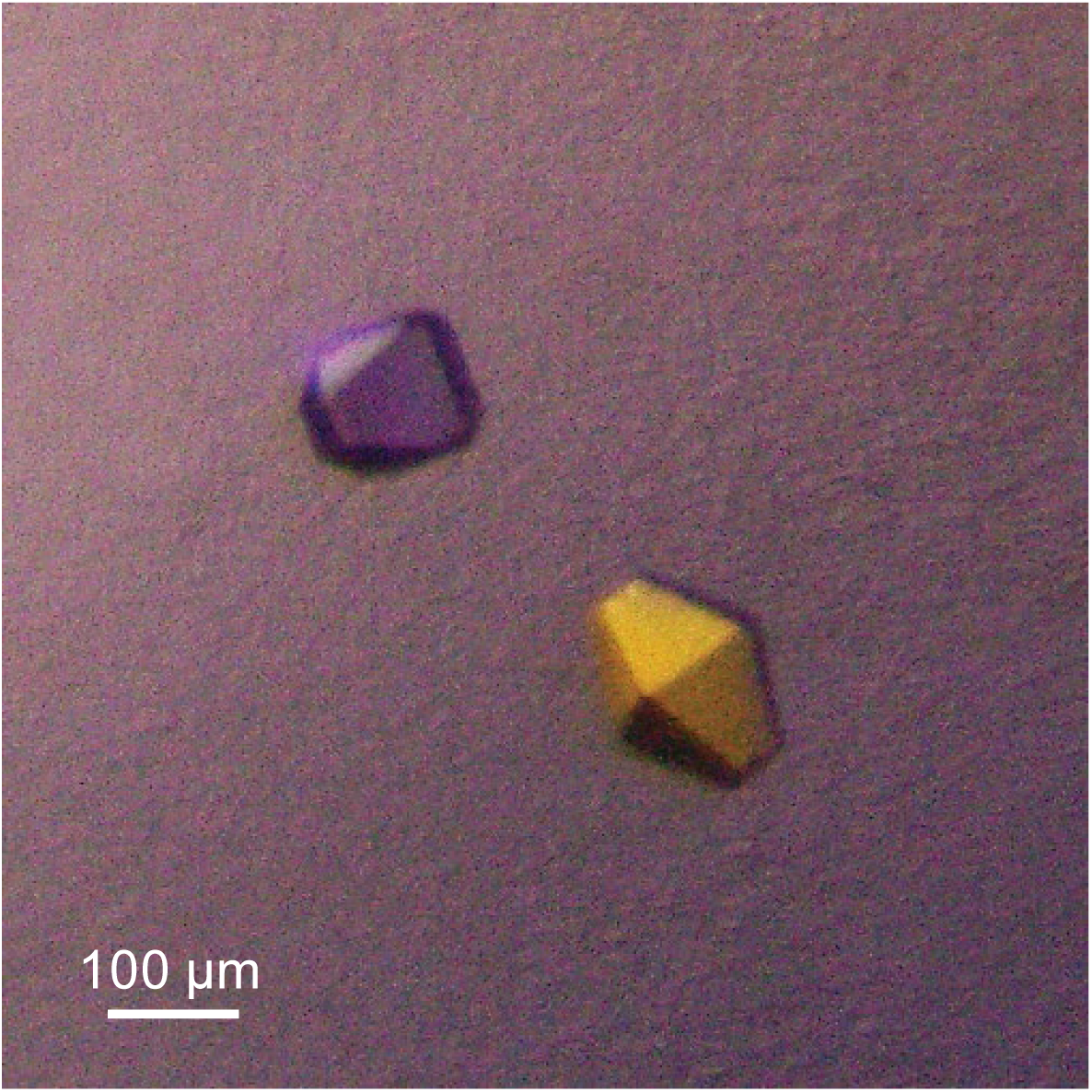
Diffraction-quality single crystals of ancD14.

**Table S1:**
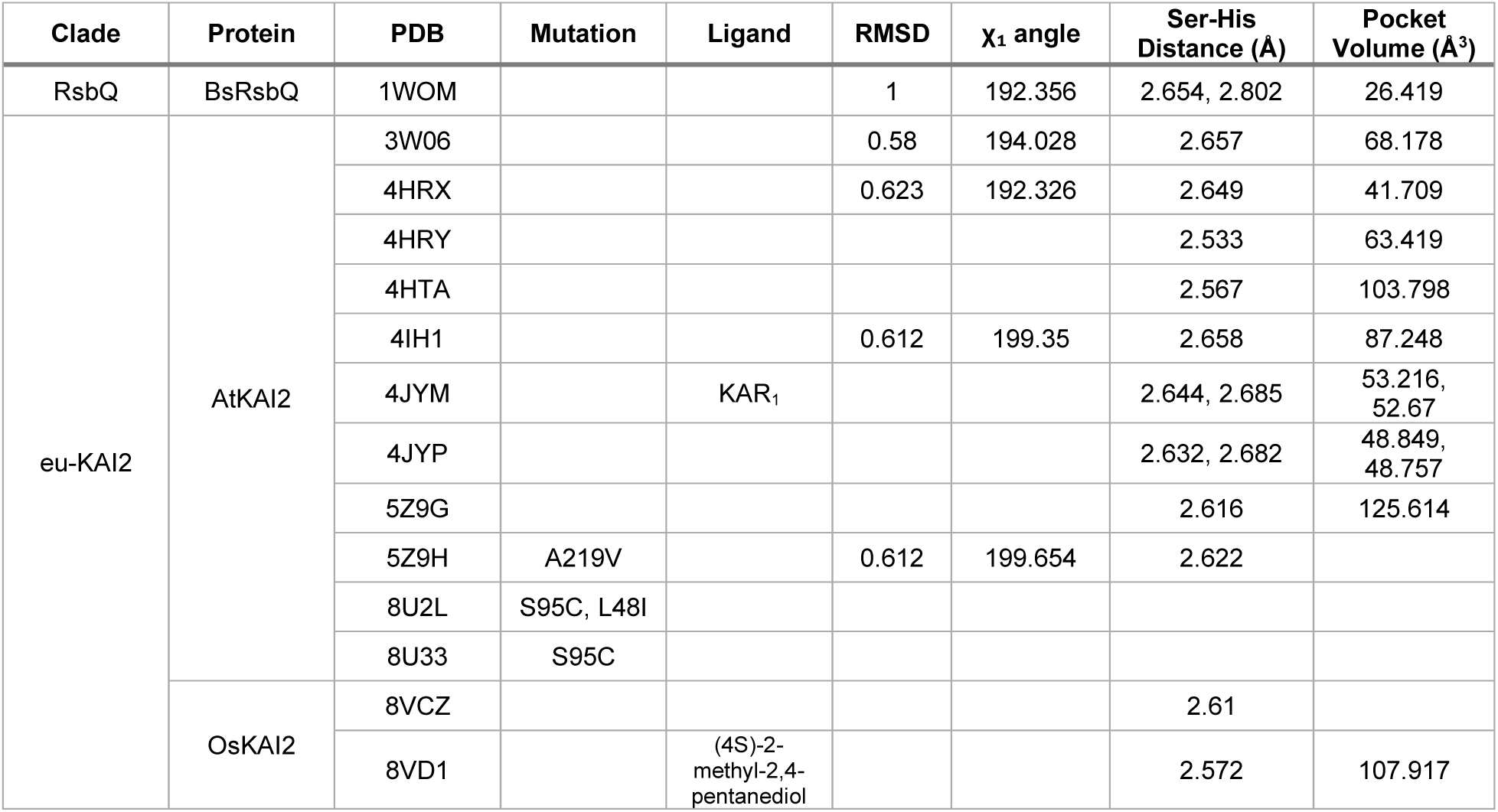
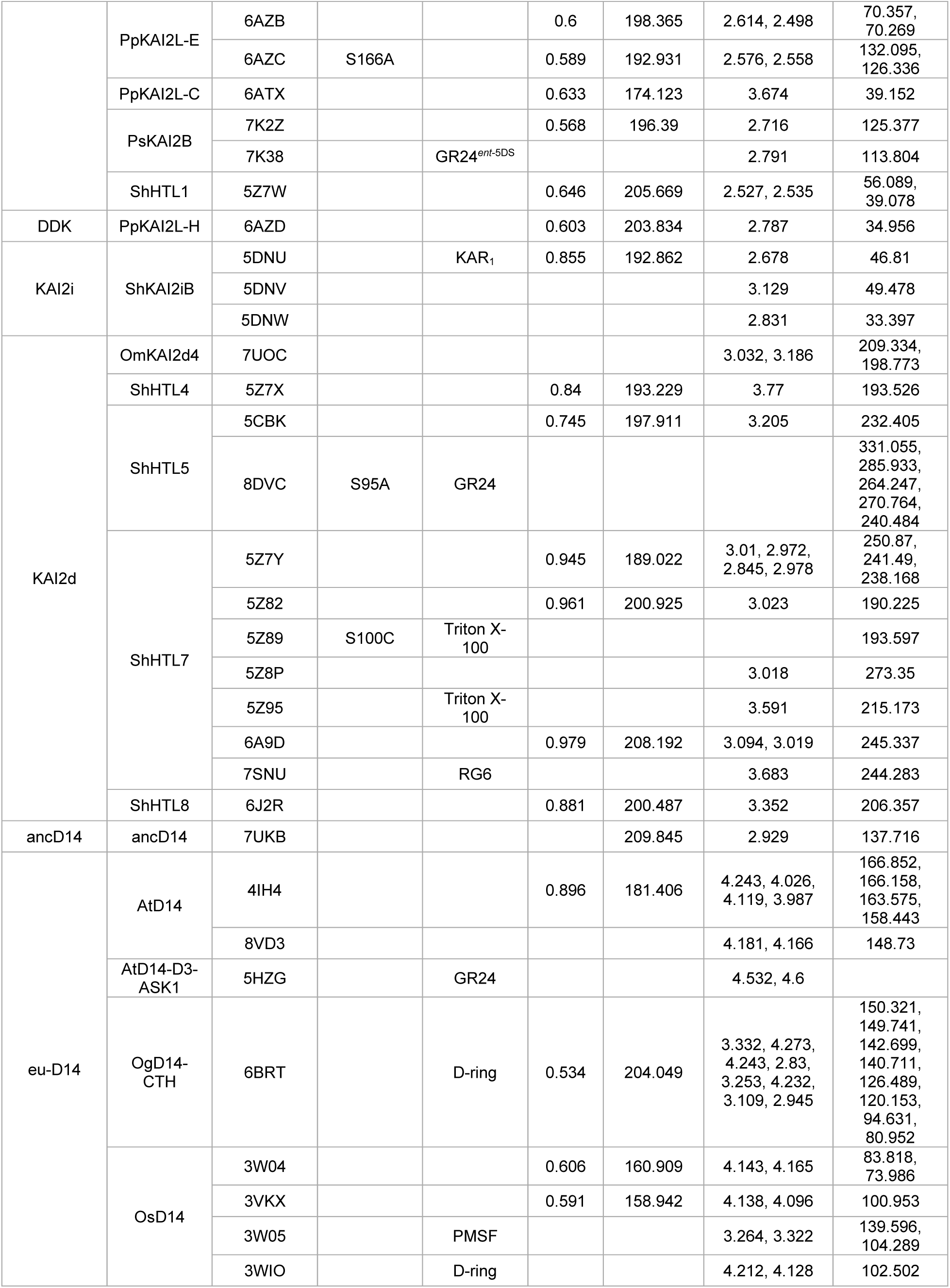
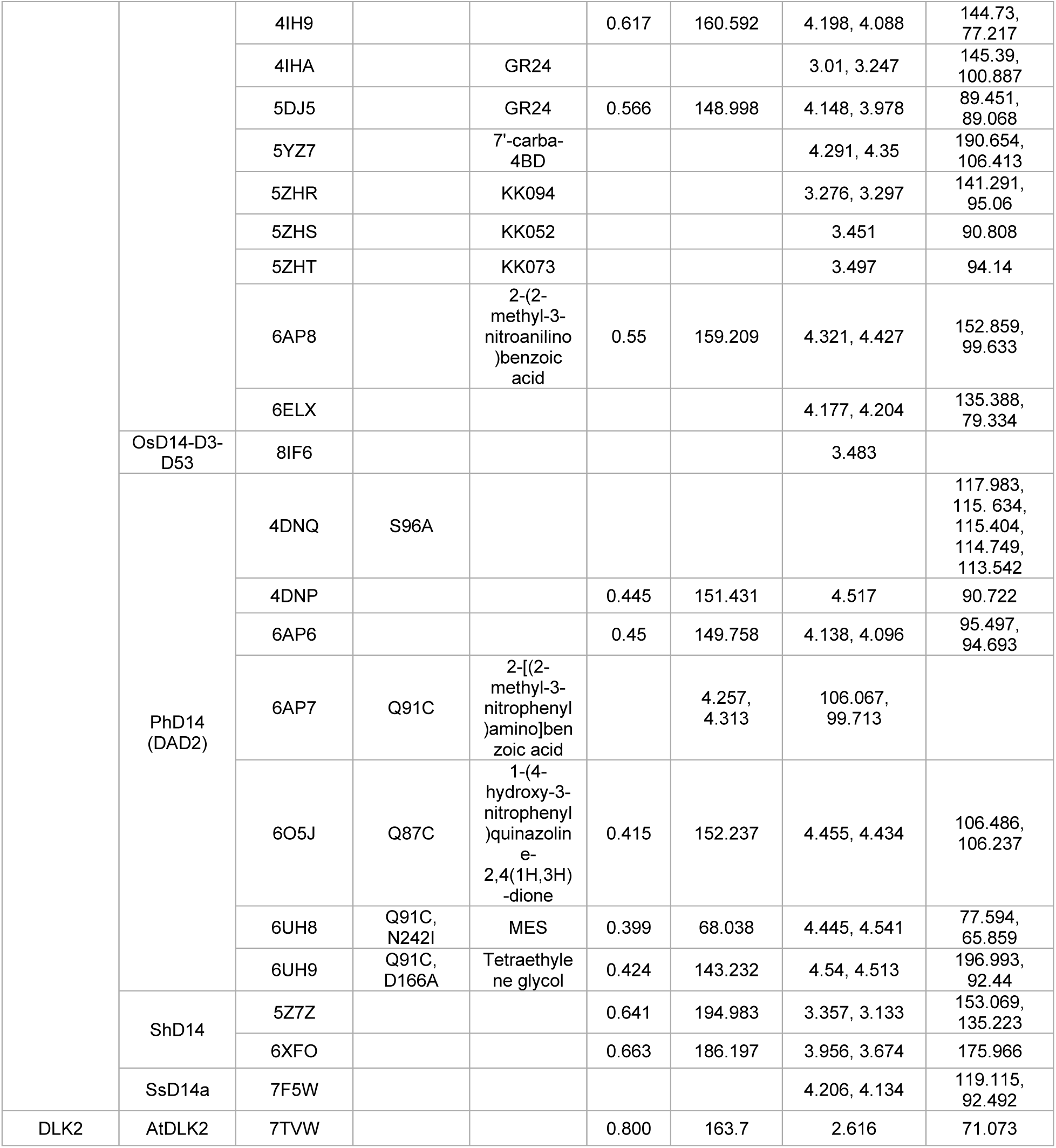
List of structures used for comparisons. Values separated by commas where measurements were made across multiple chains. RMSD refers to comparison with the ancD14 structure.

**Supplemental Data 1: Amino acid sequences used in Figure S1.**

